# Mapping-based Genome Size Estimation

**DOI:** 10.1101/607390

**Authors:** Shakunthala Natarajan, Jessica Gehrke, Boas Pucker

## Abstract

While the size of chromosomes can be measured under a microscope, obtaining the exact size of a genome remains a challenge. Biochemical methods and k-mer distribution-based approaches allow only estimations. An alternative approach to estimate the genome size based on high contiguity assemblies and read mappings is presented here. Analyses of *Arabidopsis thaliana* and *Beta vulgaris* data sets are presented to show the impact of different parameters. *Oryza sativa, Brachypodium distachyon*, *Solanum lycopersicum*, *Vitis vinifera*, and *Zea mays* were also analyzed to demonstrate the broad applicability of this approach. Further, MGSE was also used to analyze *Eschericia coli, Saccharomyces cerevisiae,* and *Caenorhabditis elegans* datasets to show its utility beyond plants. Mapping-based Genome Size Estimation (MGSE) and additional scripts are available on GitHub: https://github.com/bpucker/MGSE. MGSE predicts genome sizes based on short reads or long reads requiring a minimal coverage of 5-fold.

## Introduction

Nearly all parts of a plant are now tractable to measure, but assessing the size of a plant genome is still challenging. Although chromosome sizes can be measured under a microscope [1], the combined length of all DNA molecules in a single cell is usually unknown. About 25 years after the release of the first *Arabidopsis thaliana* genome sequence, this holds even true for one of the most important model species. Initially, biochemical methods like reassociation kinetics [2], Feulgen photometry [3], quantitative gel blot hybridization [4], southern blotting [5], and flow cytometry [6, 7] were applied. Unfortunately, these experimental methods rely on a reference genome [8]. The rise of next generation sequencing technologies [9] enabled new approaches based on k-mer profiles or the counting of unique k-mers [10, 11]. JellyFish [11], Kmergenie [12], Tallymer [13], Kmerlight [14], and genomic character estimator (gce) [15] are dedicated tools to analyze k-mers in reads. But such k-mer-based estimation methods require a high sequencing coverage [16]. Next, genome sizes can be estimated based on unique k-mers or a complete k-mer profile. Many assemblers like SOAPdenovo [17] and ALLPATHS-LG [18] perform an internal estimation of the genome size to infer an expected assembly size. Dedicated tools for genome size estimation like GenomeScope2 [19, 20] and findGSE [21] were developed. Although the authors considered and addressed a plethora of issues with real data [19], results from different sequencing data sets for the same species can vary. While some proportion of this variation can be attributed to accession-specific differences as described e.g. for *A. thaliana* [21, 22], specific properties of a sequencing library might have an impact on the estimated genome size. For example, high levels of bacterial or fungal contamination could bias the result if not removed prior to the estimation process. Due to high accuracy requirements, k-mer-based approaches are usually restricted to high quality short reads and cannot be applied to long reads of third generation sequencing technologies. The rapid development of long read sequencing technologies enables high contiguity assemblies for almost any species and is therefore becoming the standard for genome sequencing projects [23–25]. Nevertheless, some highly repetitive regions of plant genomes like nucleolus organizing region (NOR) and centromeres remain usually unassembled [22, 26, 27]. Therefore, the genome size cannot be inferred directly from the assembly size, but the assembly size can be considered a lower boundary when estimating genome sizes.

Extreme genome size estimates of *A. thaliana,* for example, 70 Mbp [2] or 211 Mbp [28], have been proven to be inaccurate based on insights from recent assemblies [22, 27, 29–32]. However, various methods still predict haploid genome sizes between 125 Mbp and 165 Mbp for *A. thaliana* accessions [29, 33–35]. Substantial technical variation is observed not only between methods, but also between different labs or instruments [36]. As described above, extreme examples for *A. thaliana* display 3 fold differences with respect to the estimated genome size. Since no assembly represents the complete genome, the true genome size remains unknown. An empirical approach, i.e. running different tools and comparing the results, might be a suitable strategy.

This work presents a conceptually different method for the estimation of genome sizes based on the mapping of reads to a high contiguity assembly. Mapping-based Genome Size Estimation (MGSE) is a Python script which processes the coverage information of a read mapping and predicts the size of the underlying genome. Since MGSE is a mapping-based method, it requires a genome sequence as reference for the read mapping process. However, this is not a limitation. The reads used for the genome size estimation, could be used for the assembly. We anticipate that future genome sequencing projects will utilize long read sequencing technologies and would easily generate assemblies of high quality which are more than appropriate for MGSE. Since MGSE relies on read mapping, it is able to support genome size estimations based on long read datasets unlike the existing kmer-based tools. Further, MGSE’s applicability to long read datasets, also helps tackle the issue of highly repetitive regions that interfere in genome size estimation. This is because long reads can span or cover the entire length of repetitive regions, ensuring correct mapping of reads and accurate coverage calculation required for an optimal genome size estimation. MGSE is an orthogonal approach to the existing tools for genome size estimation with different challenges and advantages. It is also suitable for both short and long reads obtained from different sequencing technologies, making it a broadly applicable tool in plant genomics and beyond.

## Methods

### Data sets

Sequencing data sets of the *A. thaliana* accessions Columbia-0 (Col-0) [32, 37–42] and Niederzenz-1 (Nd-1) [35] as well as several *Beta vulgaris* accessions [43–45] were retrieved from the Sequence Read Archive (AdditionalFile1). Only the paired-end fraction of the two included Nd-1 mate pair libraries was included in this analysis. Genome assembly versions TAIR9 [46], AthNd-1_v2 [27], and RefBeet v1.5 [43, 47] served as references in the read mapping process. The *A. thaliana* assemblies, TAIR9 and Ath-Nd-1_v2, already included plastome and chondrome sequences. Plastome (KR230391.1, [48]) and chondrome (BA000009.3, [49]) sequences were added to RefBeet v1.5 to allow proper placement of respective reads.

Genome sequences of *Brachypodium distachyon* strain Bd21 (GCF_000005505.3 [50]), *Solanum lycopersicum* (GCA_002954035.1 [51]), *Vitis vinifera* cultivar Chardonnay (QGNW01000001.1 [52]), *Oryza sativa* ssp. *japonica* cultivar Nipponbare (GCA_034140825.1 [53]), *Zea mays* cultivar DK105 (GCA_003709335.1 [54]), *Fragaria x ananassa* cultivar benihoppe (GCA_034370585.1 [55]), *Gossypium hirsutum* (GCF_007990345.1 [56]), *Saccharomyces cerevisiae* strain S288C (GCF_000146045.2, [57]), *Eschericia coli* strain K-12 (GCF_000005845.2, [58]), and *Caenorhabditis elegans* strain Bristol N2 (GCF_000002985.6, [59]) were retrieved from the NCBI. Corresponding read data sets of *Brachypodium distachyon* ([50]), *Solanum lycopersicum* [51, 60–62], *Vitis vinifera* [52], *Oryza sativa* [53, 63], *Zea mays* ([54]), *Fragaria x ananassa*, *Gossypium hirsutum*, *Saccharomyces cerevisiae* [64], *Eschericia coli* [65], and *Caenorhabditis elegans* [66] were retrieved from the Sequence Read Archive (AdditionalFile1). These read datasets were chosen based on similarly analyzed material to the respective reference genome sequence. This was done to ensure that the genome size estimation is representative for the reference strain and is not deviating due to structural differences between different strains.

### Genome size estimation

JellyFish2 v2.2.4 [11] was applied for the generation of k-mer profiles which were subjected to GenomeScope2 [19]. Selected k-mer sizes ranged from 19 to 25. Results of different sequencing data sets and different k-mer sizes per accession were compared. Genomic character estimator (gce) [15] and findGSE [21] were applied to infer genome sizes from the k-mer histograms. If tools failed to predict a value or if the prediction was extremely unlikely, values were masked to allow meaningful comparison and accommodation in one figure. The number of displayed data points (i.e. successful predictions) is consequently a quality indicator.

### Mapping-based genome size estimation

Despite some known biases [67–69], the underlying assumption of MGSE is a nearly random fragmentation of the DNA and thus an equal distribution of sequencing reads over the complete sequence. If the sequencing coverage per position (C) is known, the genome size (N) can be calculated by dividing the total amount of sequenced bases (L) by the average coverage value: N = L / C. The working concept of MGSE is explained in Fig. 1.

**Figure 1:**
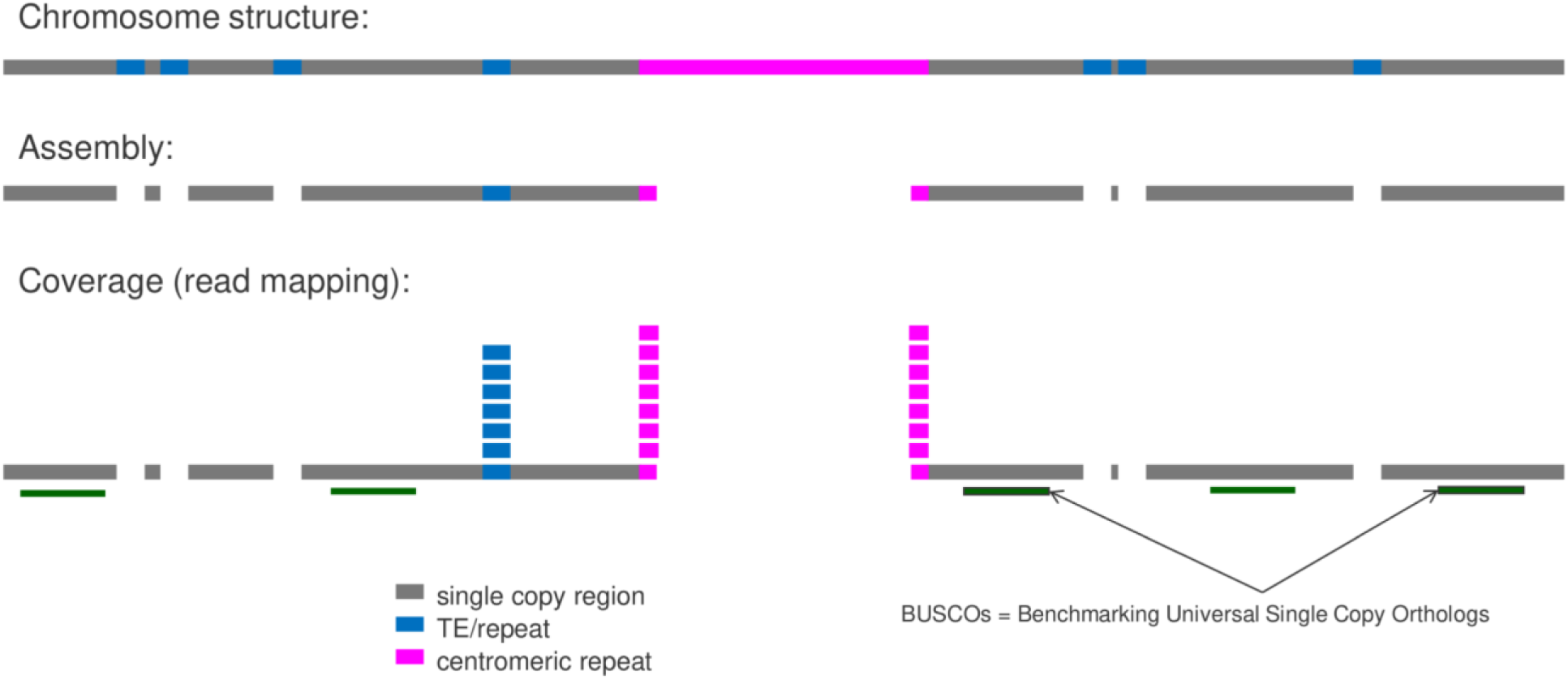
Concept diagram showing the working logic behind read mapping-based genome size estimation. An assembly is required as a basis for the analysis, but does not have to be perfect.

Underrepresented repeats and other regions like the unresolved paralogs of the SEC10 gene in the *Arabidopsis thaliana* genome sequence [70], display a higher coverage, because reads originating from different genomic positions are mapped to the same sequence. The accurate identification of the average coverage is crucial for a precise genome size calculation. Chloroplastic and mitochondrial sequences account for a substantial proportion of reads in sequencing data sets, while contributing very little size compared to the nucleome. Therefore, sequences with very high coverage value, i.e., plastome and chondrome sequences are included during the mapping phase to allow correct placement of reads, but can be excluded from MGSE in a later stage. A user-provided list of reference regions is used to calculate the median or mean coverage based on all positions in these specified regions. Benchmarking Universal Single Copy Orthologs (BUSCO) [71, 72] can be deployed to identify such a set of *bona fide* single copy genes which should serve as suitable regions for the average coverage calculation. Since BUSCO is frequently applied to assess the completeness of a genome assembly, these files might be already available to users. GFF3 files generated by BUSCO can be concatenated and subjected to MGSE. As some BUSCOs might occur with more than one copy, MGSE provides an option to reduce the predicted gene set to the actual single copy genes among all identified BUSCOs.

For short reads, BWA MEM v0.7 [73] was applied for the read mapping and MarkDuplicates (Picard tools v2.14) [74] was used to filter out reads originating from PCR duplicates. For long reads, minimap2 v2.24 [75, 76] was used for read mapping. Next, a previously described Python script [77] was deployed to generate coverage files from the BAM files, which provides information about the number of aligned sequencing reads covering each position of the reference sequence. This process involves bedtools v2.30.0 [78]. Finally, MGSE v3.1 (https://github.com/bpucker/MGSE) was run on these coverage files to predict genome sizes independently for each data set.

### Coverage threshold analysis

MGSE relies on coverage calculation for estimating the genome sizes. Hence, it is important to know the minimum coverage that datasets given to MGSE must have for getting optimal results. For this, all the long and short read datasets of *Arabidopsis thaliana*, described earlier, were taken and sub-sampled into FASTQ files containing varying percentages of the reads - 100%, 75%, 50%, 25%, 10%, 7.5%, 5%, 2.5%, 1%, and 0.5% using seqtk [79]. Then each of these different read files across all the samples were given to MGSE to obtain genome size estimates. The number of sequenced bases in the read files given to MGSE and the genome size estimates of MGSE were correlated to determine the minimum coverage of long and short read datasets that MGSE can process. The above sub-sampled files were also given to GenomeScope2 to obtain a comparative coverage threshold value for short read datasets that can be handled by this kmer-based tool.

### Runtime analysis

Runtime is an important factor when users determine the resources needed for using a particular tool. Here, runtime analyses were conducted by correlating the number of sequenced bases in the read files given to MGSE and the total time taken for a complete run including indexing, read mapping, coverage calculation, and the final genome size prediction. Since minimap2 is slightly more efficient than BWA-MEM, some long read datasets were subjected to MGSE runs with a single CPU. Thereafter, the number of CPUs for the MGSE runs on short and long reads was set to be 10, to evaluate the runtimes with moderate resources. Since read mapping is already a part of most genome assembly processes, users might already have BAM files at hand that could serve as input for MGSE. Therefore, another set of runtime analyses were conducted by correlating the size of the input datasets and only the times taken for coverage calculation and genome size prediction. These MGSE runtimes were then compared with GenomeScope2 runtimes for the same datasets carried out with 10 CPUs to benchmark and understand the performance of MGSE in direct comparison to a kmer-based tool.

## Results & discussion

### *Arabidopsis thaliana* genome size

MGSE was deployed to calculate the genome size of the two *A. thaliana* accessions Col-0 and Nd-1 (Fig. 2). In order to identify the best reference region set for the average coverage calculation, different reference region sets were tested. Manually selected single copy genes, all protein encoding genes, all protein encoding genes without transposable element related genes, only exons of these gene groups, and BUSCOs were evaluated (AdditionalFile2). The results were compared against predictions from GenomeScope2, gce, and findGSE for k-mer sizes 19, 21, 23, and 25.

**Fig. 2:**
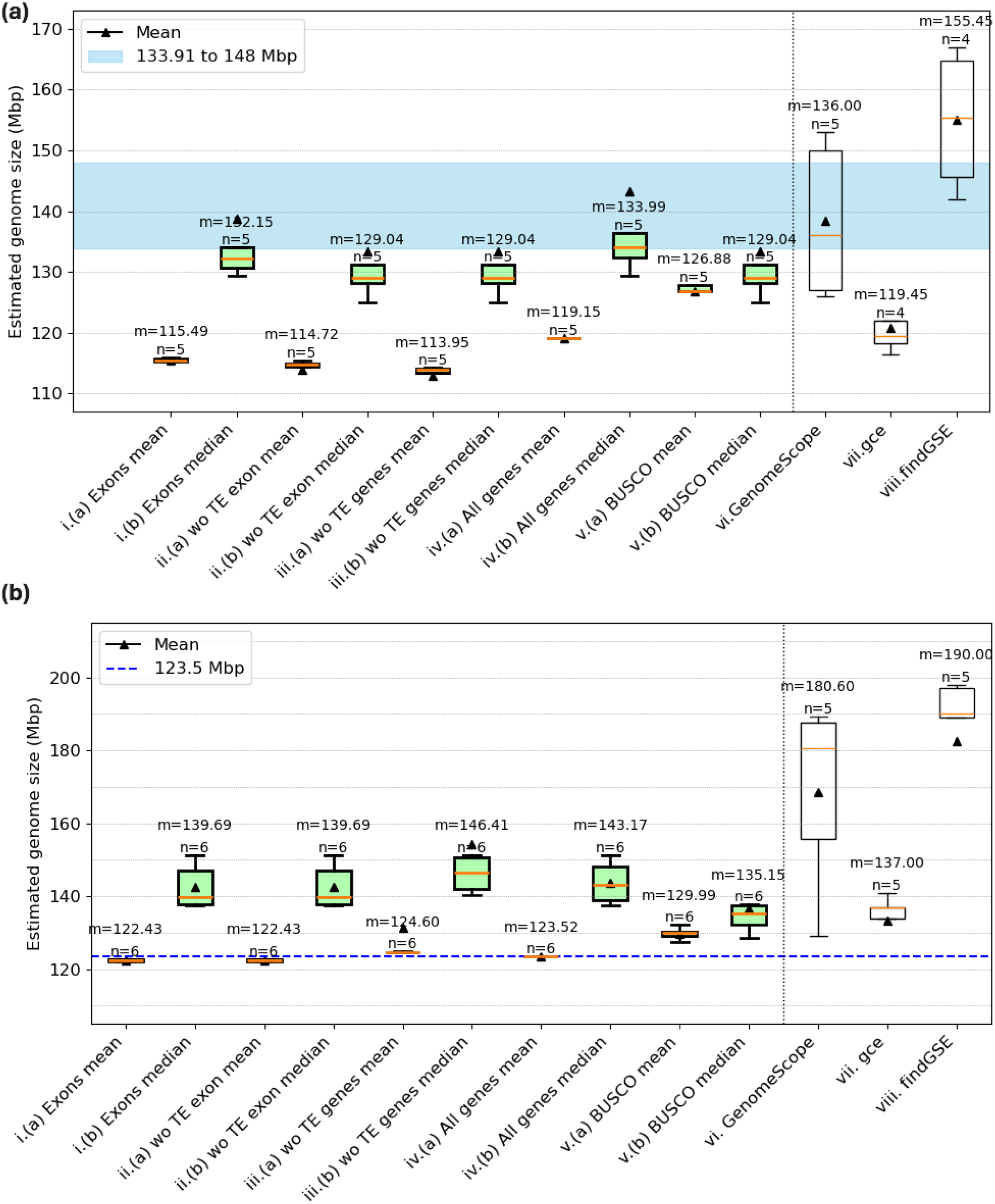
Comparison of *Arabidopsis thaliana* genome size estimations. Genome sizes of the *A. thaliana* accessions Col-0 (a) and Nd-1 (b) were predicted by MGSE, GenomeScope2, gce, and findGSE. Different MGSE approaches were evaluated, differing by the set of regions for the average coverage calculation (e.g. all genes) and the methods for the calculation of this value (mean/median). Multiple read data sets (n) were analyzed by each tool/approach to infer an average genome size given as median (m, yellow line) and mean (black triangles), transposable elements = TE, without = wo. The blue region in (a) shows the expected genome size range. It has the near complete assembly size of Col-0 [80] as the lower boundary and one of the largest reported assembly sizes of *Arabidopsis thaliana* [81] as the upper boundary (AdditionalFile8); The blue line in (b) shows a high quality and the largest reported assembly size for Nd-1 till date.

Many estimation approaches predict a genome size for Col-0 that is below the largest reported assembly size of 148 Mbp for *Arabidopsis thaliana* [81] and display substantial variation between samples. The BUSCO-based approaches with MGSE appeared promising due to low variation between different samples and a prediction close to the average sizes of almost complete Col-0 genome sequences. GenomeScope2 predicted a slightly higher genome size with greater variation between samples, while gce reported consistently much smaller values. findGSE predicted on average a substantially larger genome size. Finally, sample sizes below five (for Col-0) and below six (for Nd-1) indicated that prediction processes failed for individual approaches e.g. due to insufficient read numbers.

Due to low variation between different samples and a plausible average genome size, the BUSCO-based approaches appeared promising for Nd-1 as well (Fig. 2b). The reported assembly size of 123.5 Mbp is known to be an underestimation of the true genome size of the Nd-1 accession [27]. The genome size estimation of about 135 Mbp inferred for Nd-1 through the BUSCO approach of MGSE is also slightly below previous estimations of about 146 Mbp [35]. Approximately 123.5 Mbp are assembled into pseudochromosomes which do not contain complete NORs or centromeric regions [27]. Based on the read coverage of the assembled 45S rDNA units, the NORs of Nd-1 are expected to account for approximately 2-4 Mbp [35].

Centromeric repeats which are only partially represented in the genome assembly [27] account for up to 11 Mbp [35]. In summary, the Nd-1 genome size is expected to be around 138-140 Mbp. The single-copy BUSCO genes emerged as the best set of reference regions for MGSE based on further analyses using the Ath-Nd1_v2 assembly. In summary, MGSE (considering the BUSCO median estimation) and gce predicted reasonable genome sizes for Nd-1. The average predictions by GenomeScope2 and findGSE are very unlikely, as they contradict most estimations of *A. thaliana* genome sizes [6, 21, 27, 35].

MGSE was also used to estimate the genome size of the Col-0 accession using long read data sets of 14 GABI-Kat lines [32] (Fig. 3, AdditionalFile3). Since estimates relying on single copy BUSCO genes as reference regions were identified as the best approach in previous *A. thaliana* analyses, the same strategy was applied to datasets of all these GABI-Kat lines.

**Fig. 3:**
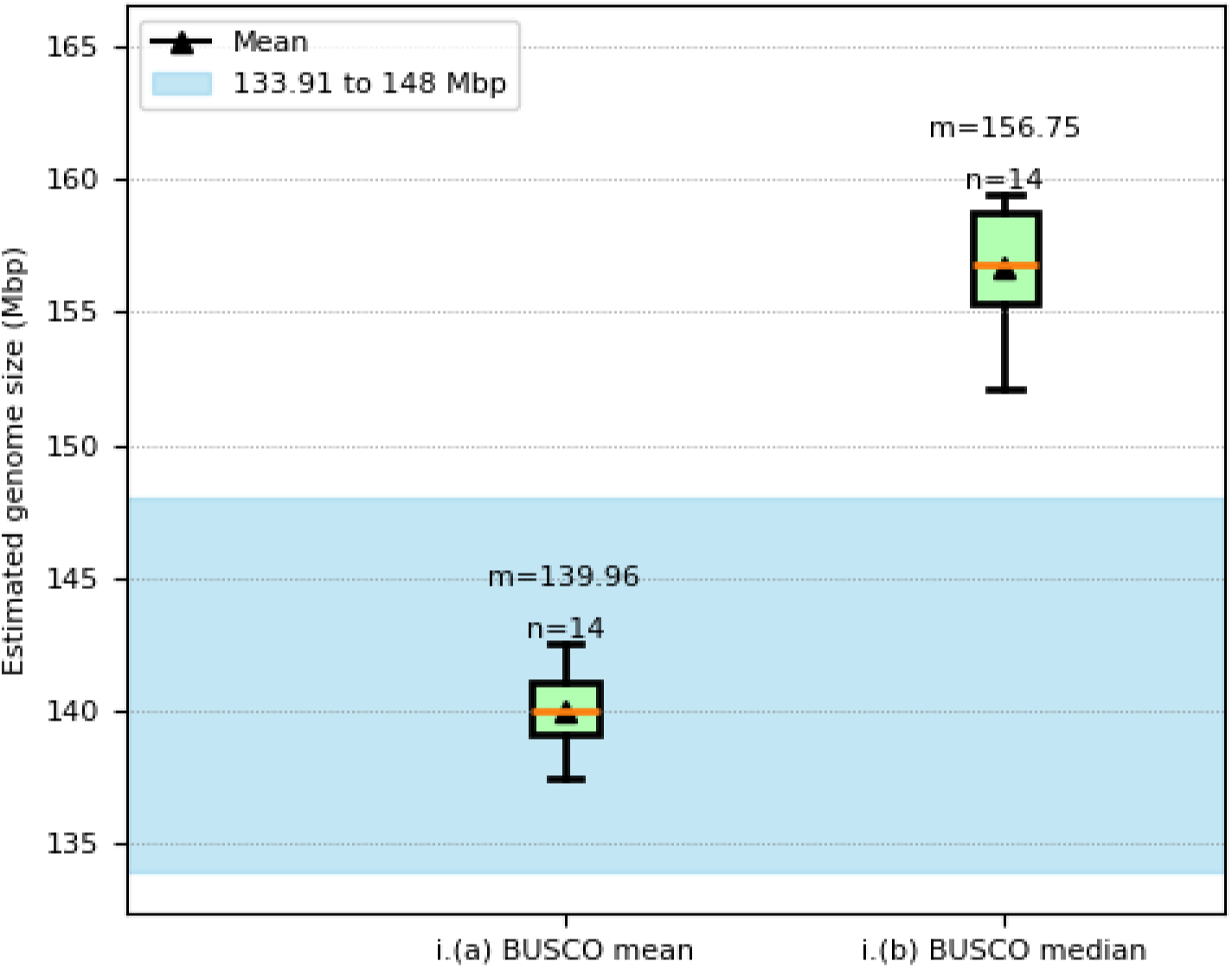
MGSE estimations using long read sequencing data sets of 14 GABI-Kat lines (Col-0 accession). The blue region shows the expected genome size range with the near complete assembly size of Col-0 [80] as the lower boundary and one of the largest reported assembly sizes of *Arabidopsis thaliana* [81] as the upper boundary (AdditionalFile8).

The MGSE BUSCO mean fits well into the expected genome size range of Col-0, while the median estimate is slightly higher than the largest reported assembly size considered here. This deviation could be attributed to large numbers of InDels in long reads. There are a number of methods to reduce this bias in the input long read data, like the use of PacBio HiFi technology [82] and application of HERRO-correction to ONT reads [83], which could help improve the MGSE estimates. Nevertheless, the estimates still fall into the acceptable limits for *Arabidopsis* genome size estimates. This depicts that MGSE is able to process long read datasets unlike the kmer-based approaches, making it valuable as long read sequencing is gaining importance.

The feasibility of MGSE was further demonstrated by estimating the genome sizes of 1,028 *A. thaliana* accessions (Fig. 4, AdditionalFile4) which were analyzed by re-sequencing as part of the 1001 genome project [84]. Most predictions by MGSE v0.55 are between 120 Mbp and 160 Mbp, while all other tools predict most genome sizes between 120 Mbp and 200 Mbp with some outliers showing very small or very large genome sizes. MGSE differs from all three tools when it comes to the number of failed or extremely low genome size predictions. All k-mer-based approaches predicted genome sizes below 50 Mbp, which are most likely artifacts, possibly due to very low sequencing coverage. This comparison revealed systematic differences between findGSE, gce, and GenomeScope2 with respect to the average predicted genome size. findGSE tends to predict larger genome sizes than gce and GenomeScope2. Very large genome sizes could have biological explanations like polyploidization events.

**Fig. 4:**
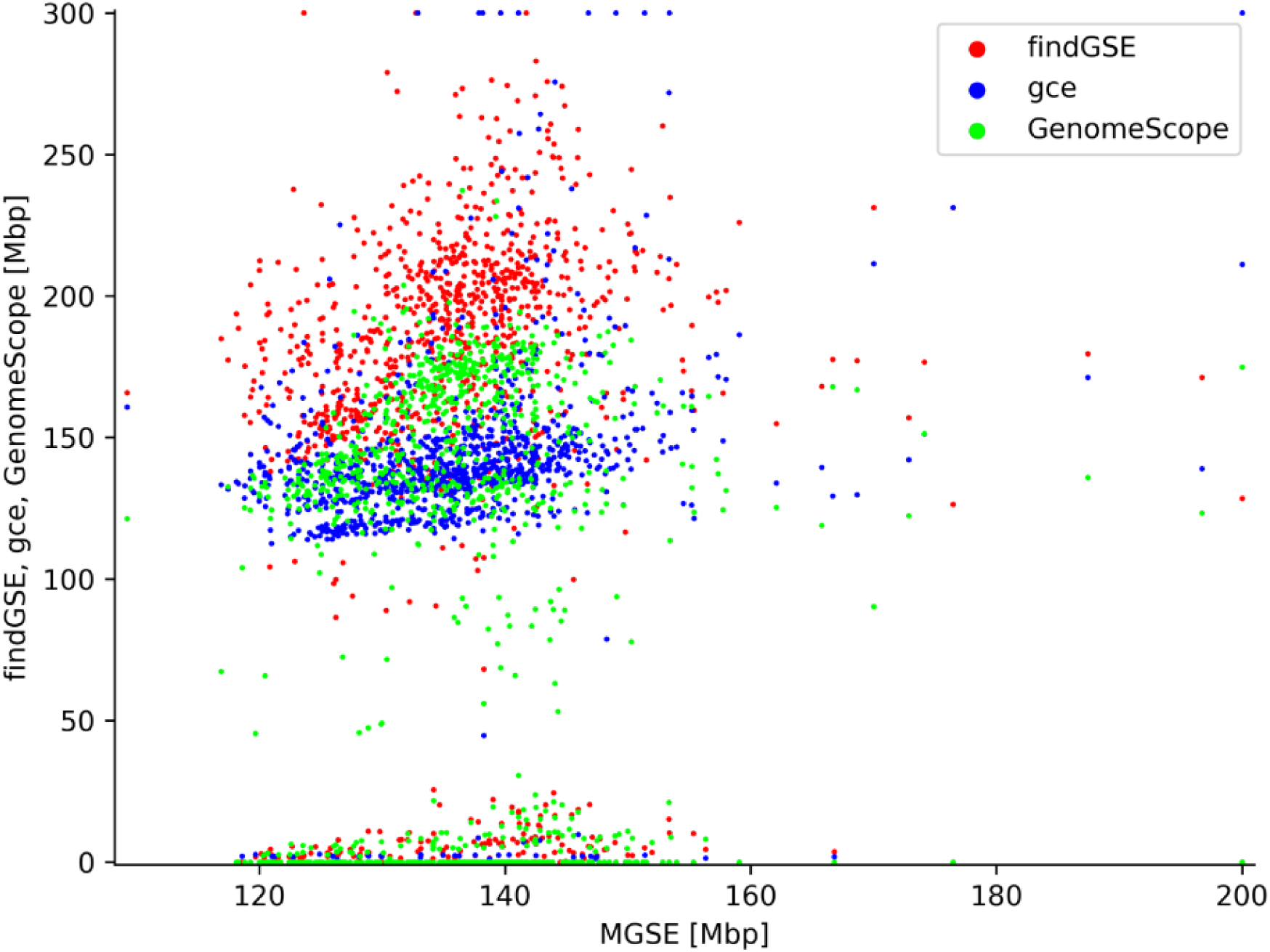
Genome size estimations of *Arabidopsis thaliana* accessions. MGSE, findGSE, gce, and GenomeScope2 were deployed to predict the genome sizes of 1,028 *A. thaliana* accessions based on sequence read data sets (AdditionalFile4). Extreme outliers above 200 Mbp (MGSE) or 300 Mbp (other tools) are displayed at the plot edge to allow accommodation of all data points with sufficient resolution in the center.

### *Beta vulgaris* genome size

Different sequencing data sets of *Beta vulgaris* were analyzed via MGSE, GenomeScope2, gce, and findGSE to assess the applicability to larger and more complex genomes (Fig. 5, AdditionalFile5). Different cultivars served as material source for the generation of the analyzed read data sets. Therefore, minor differences in the true genome size are expected. Moreover, sequence differences like single nucleotide variants, small insertions and deletions, as well as larger rearrangements could influence the outcome of this analysis. The reference sequence RefBeet v1.5 already represented 567 Mbp [43, 47], but an investigation of the repeat content indicates a larger genome size due to a high number of repeats that are not represented in the assembly [85]. The largest reported assembly of the reference accession KWS2320 has a size of 573 Mbp [86] thus all estimations below 573 Mbp can be considered as underestimations. Given that the assembly size of the accession U2Bv is even slightly larger with 596 Mbp [86]. When assuming centromere sizes of only 2-3 Mbp per chromosome, the assembly size of 600 Mbp could be close to the true genome size. Previous genome size estimations reported 714-758 Mbp [6] and 731 Mbp [43]. In summary, it appears likely that the true genome size of sugar beet is between about 600 Mbp and 758 Mbp.

**Fig. 5:**
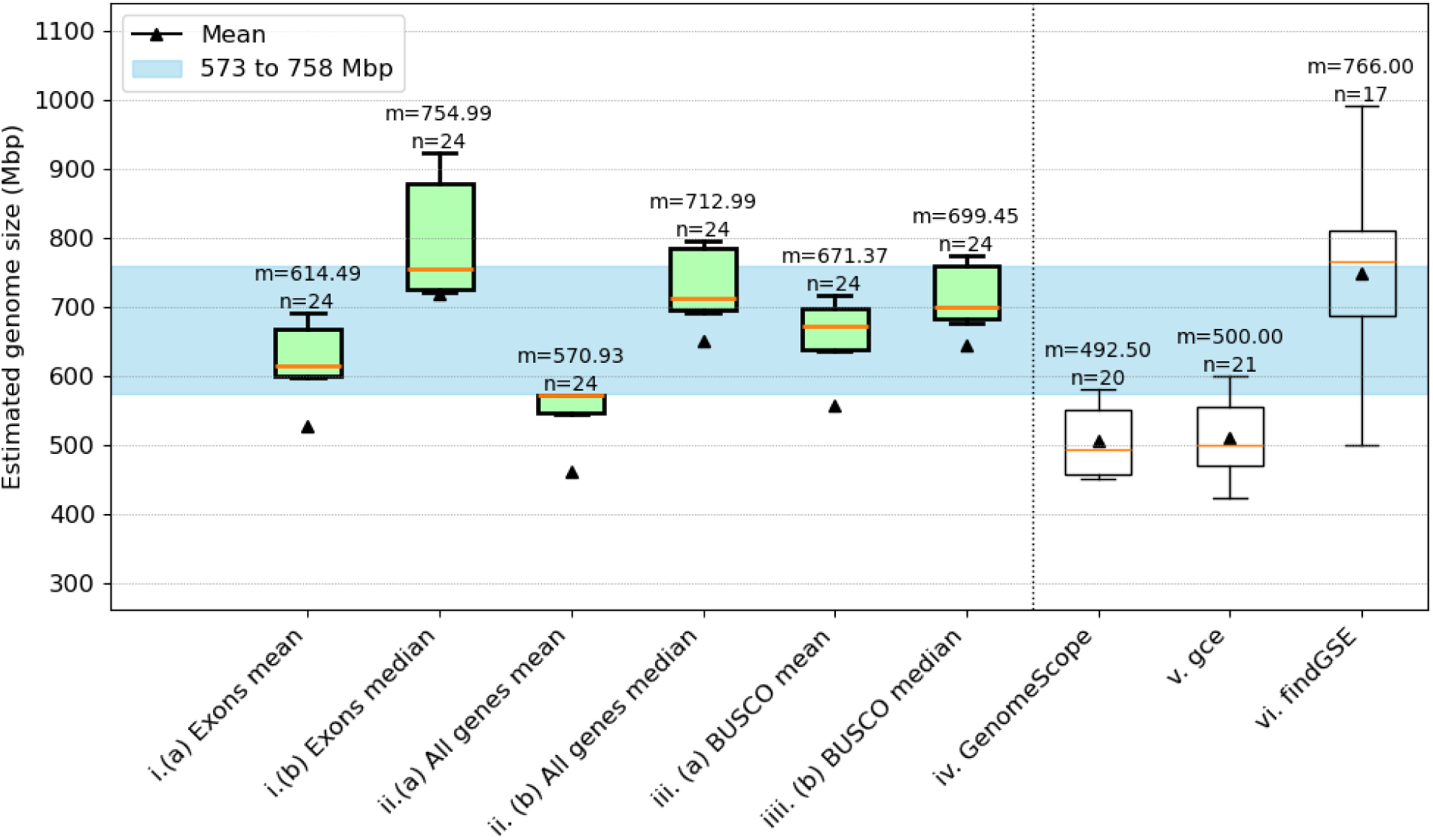
Comparison of *Beta vulgaris* genome size estimations. The genome size of *B. vulgaris* was predicted by MGSE, GenomeScope2, gce, and findGSE. Different MGSE approaches were evaluated differing by the set of regions for the average coverage calculation (e.g. all genes) and the methods for the calculation of this value (mean/median). Multiple read data sets (n) were analyzed by each tool and approach to infer an average genome size given as median (m, yellow line) and mean (black triangles). The blue region shows the expected genome size range with the largest reported assembly for *Beta vulgaris* [86] as the lower boundary and the largest previously estimated genome size of *Beta vulgaris* [6] as the upper boundary (AdditionalFile8).

In this study, a few samples were considered to be outliers as these samples showed low coverage values. It is possible that these samples belong to a different subspecies which could explain a low mapping rate against the sugar beet reference. Nevertheless, MGSE, and findGSE performed best in estimating the genome sizes of *B. vulgaris*. findGSE estimated extremely variable values, but mostly around the previously estimated genome sizes [6, 43]. It is noticeable that the findGSE analysis failed for a number of samples. It is also important to note that GenomeScope2 and gce underestimate the genome size. The prediction of less than 600 Mbp is an underestimation, because this value is below the size of available genome sequences. Looking at different MGSE modes, the non-BUSCO approaches gave extremely high estimates and showed high variability between samples, the BUSCO-based approaches were in a plausible range. Therefore, the mean and median-based approaches relying on all genes or just the BUSCO genes as reference regions for the sequencing coverage estimation outperformed all other approaches. When comparing the *A. thaliana* and *B. vulgaris* analyses, the calculation of an average coverage in all single-copy BUSCO genes appears to be the most promising approach. This aligns well with the assumption that single copy BUSCO genes should appear with exactly one copy in the genome and would be prime regions to infer the average sequencing coverage depth.

### *Oryza sativa* genome size

Rice (*Oryza sativa*) is a major food grain and belongs to the monocot lineage. Given its importance as a food crop, there have been a lot of efforts dedicated to obtaining a complete genome sequence of rice [87]. Recently, a complete genome sequence of *Oryza sativa* ssp. japonica cv. Nipponbare was reported with 385.7 Mbp [53]. It was obtained using a hybrid assembly approach combining reads from Illumina, ONT, and PacBio sequencing technologies with a Hi-C dataset [53].

MGSE was used to estimate the size of the rice genome using reads from different sequencing technologies (Illumina, ONT, and PacBio). Other tools like findGSE, gce, and GenomeScope2 were also used to estimate the genome size using the Illumina reads. As these tools are relying on k-mers, they have only been recommended for the use with short reads. The estimation results using rice Illumina reads are shown in Fig. 6a. The comparative estimations by MGSE using short and long reads are shown in Fig. 6b.

**Fig. 6:**
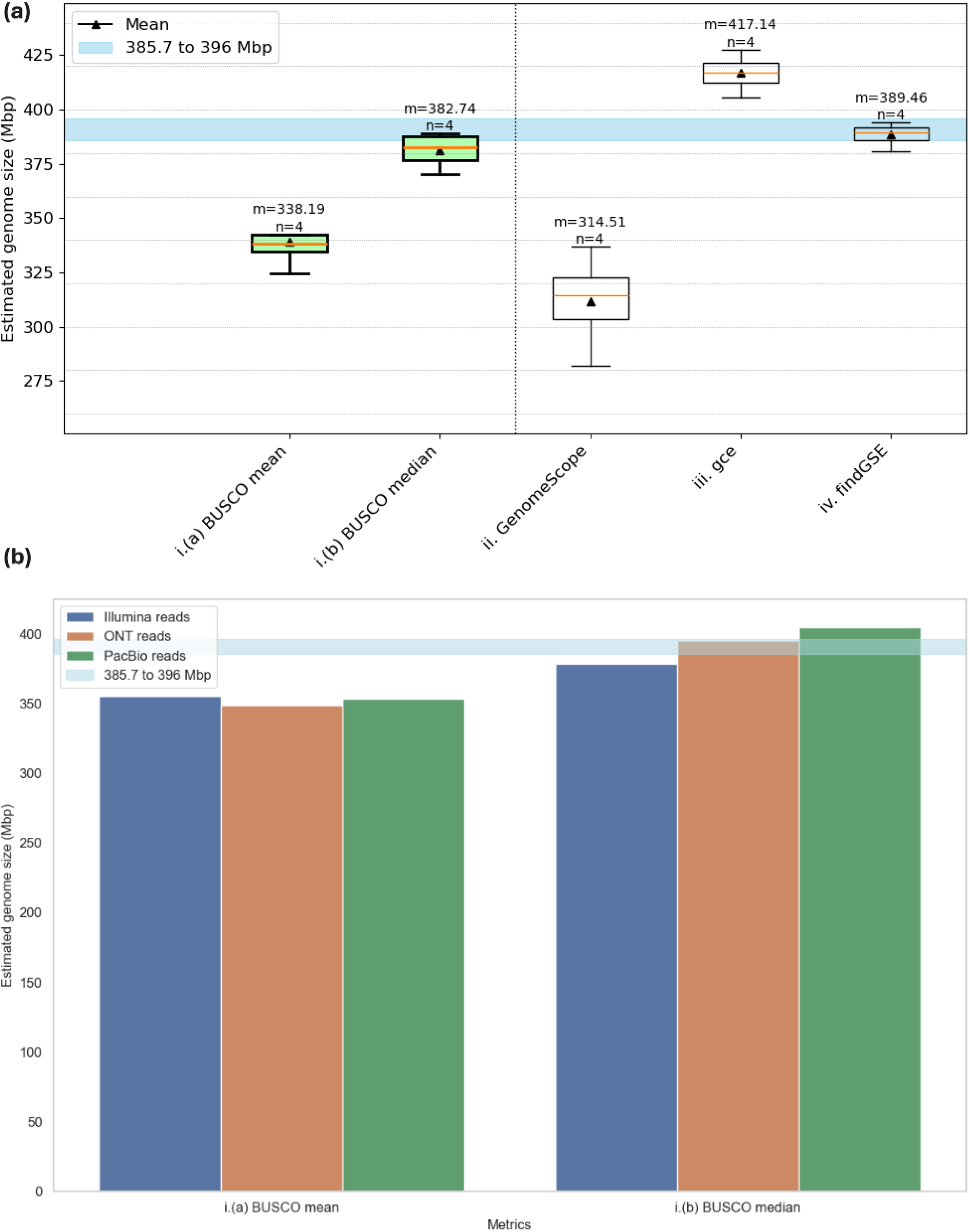
(a) Comparison of *Oryza sativa* genome size estimations using short reads. (b) Comparison of MGSE genome size estimations on short and long reads of *O. sativa.* The blue region shows the expected genome size values (AdditionalFile8). It has the size of the near complete genome sequence of *O. sativa* as the lower boundary and the size of the largest reported assembly of *O. sativa* as the upper boundary [88].

While MGSE BUSCO median and findGSE estimate sizes very close to the size of the presumable complete genome sequence, BUSCO mean estimates of MGSE and GenomeScope2 underestimate the genome size. gce slightly overestimates the size.

In contrast to the other tools, MGSE can perform genome size predictions based on long reads. In fact, BUSCO median estimates with MGSE are very close to the expected genome size and they appear to be even more plausible than the estimations based on short reads.

### Application to broad taxonomic range of species

After optimization of MGSE on *A. thaliana* (Rosids) and *B. vulgaris* (Caryophyllales), the tool was deployed to analyze data sets of different taxonomic groups thus demonstrating broad applicability. *Brachypodium distachyon* was selected as representative of grasses, *Solanum lycopersicum* represents the Asterids, *Zea mays* was included as monocot species with high transposable element content in the genome, *Vitis vinifera* was selected due to a very high heterozygosity, *Fragaria x ananassa, Gossypium hirsutum* were added to represent polyploid species, *Eschericia coli, Saccharomyces cerevisiae, and Caenorhabditis elegans* were chosen to represent bacterial, fungal and animal genomes. The predictions of MGSE are generally in the same range as the predictions generated by GenomeScope2, gce, and findGSE (AdditionalFile5, AdditionalFile9, AdditionalFile10, AdditionalFile11, and AdditionalFile12). With an average prediction of 290 Mbp as genome size of *B. distachyon*, the MGSE prediction is slightly exceeding the assembly size. findGSE prediction also slightly exceeds the assembly size. GenomeScope2 and gce predict genome sizes below the assembly size (AdditionalFile7). The *Z. mays* genome size is underestimated by all four tools. However, MGSE outperforms GenomeScope2 and gce on the analyzed data sets (AdditionalFile8). The *S. lycopersicum* genome size is underestimated by MGSE on most data sets. However, the compared tools failed to predict a genome size for multiple read data sets (AdditionalFile9). MGSE estimates for *V. vinifera* were well within the expected genome size range. findGSE and gce overestimated the genome size. GenomeScope2 underestimated the genome size of *V. vinifera* as evident from the assembly size exceeding the estimation (AdditionalFile10).

To further assess MGSE’s applicability to complex polyploid genomes, MGSE was deployed on *Fragaria x ananassa* and *Gossypium hirsutum*. *Fragaria x ananassa* was selected because it is a plant with high ploidy (octoploid) and is also an important crop with global relevance. From the MGSE results, it is evident that, selecting all the genes as well as BUSCO genes, as reference genes for coverage calculation gives optimal results for *Fragaria ananassa*, albeit with a slight overestimation of the median genome size (AdditionalFile13). But it is important to note that polyploid genomes have multiple copies of a region in different subgenomes, resulting in very high coverage values. Hence, it is recommended to select the ‘--ignore’ option to deactivate the blacklisting of high coverage contigs in the case of polyploids. Next, *Gossypium hirsutum*, was selected, as it is an allotetraploid plant with a high degree of repeats and is also an important crop. While MGSE performs optimally when choosing all the genes for coverage calculation and choosing the ‘--ignore’ option for *Gossypium hirsutum*, it fails to predict plausible genome sizes in the other cases (AdditionalFile14). This could be attributed to the high degree of repetitive genomic sequences in upland cotton [89, 90]. Hence, it is suggested to use MGSE with the ‘--ignore’ option, while deploying it on polyploid genomes, especially those with a high number of repeat elements.

Further, it is necessary to note here that MGSE predicts the genome size of the organism, based on the ploidy of the genome assembly provided by the user. To help the user assess the ploidy level of the assembly that they are providing and to facilitate the selection of optimal parameters for running MGSE, we developed an additional script. ‘busco2ploidy.py’ that estimates the assembly’s ploidy by analyzing BUSCO results (i.e. BUSCO gene duplications) of the assembly. Alternatively, we recommend using tools like smudgeplot [20] which help infer the ploidy before deciding upon the best options for an MGSE run.

Moreover, MGSE is suitable for organisms across a wide taxonomic range as depicted for bacterial (*Eschericia coli*), fungal (*Saccharomyces cerevisiae*) and animal (*Caenorhabditis elegans*) genomes. MGSE estimates for *E. coli* and *C. elegans* aligned well with the expected genome sizes of these organisms (Fig. 7a, 7c). But, MGSE slightly overestimates the yeast genome size when compared to the standard reference assembly size (Fig. 7b). However, there are some studies that reported yeast strains with additional genomic regions compared to the standard reference assembly [91]. Further, some recent studies also report a few genome assembly sizes of yeast strains to exceed the value of 12.1 Mb reported for the standard reference [92], corroborating the slightly higher MGSE estimates for yeast.

**Fig. 7:**
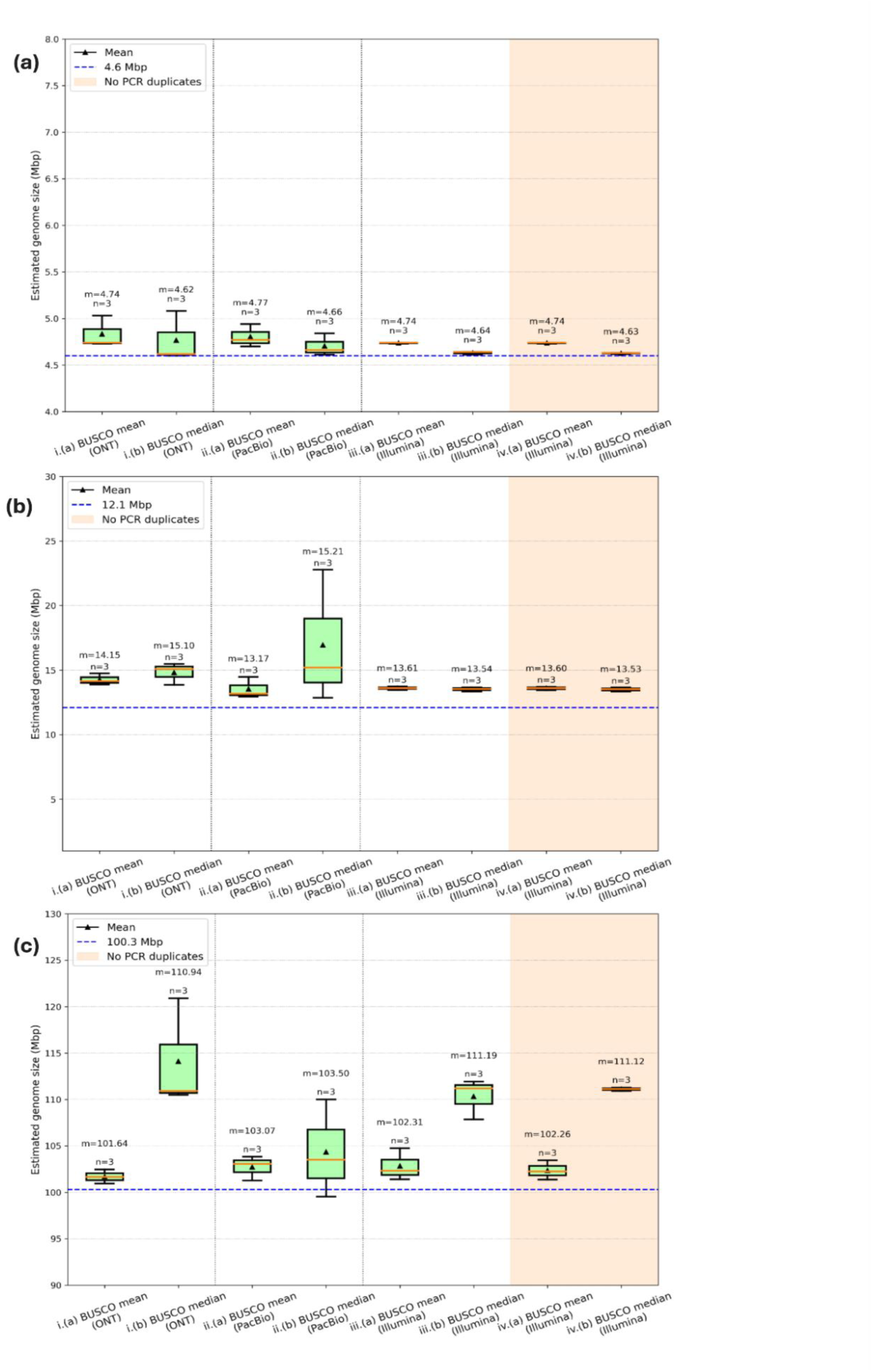
MGSE genome size estimates of (a) *Eschericia coli* (b) *Saccharomyces cerevisiae* (c) *Caenorhabditis elegans.* The dotted blue lines denote the genome assembly sizes of the standard reference organisms. The y-axis was automatically set to a range that highlights differences between the genome size prediction approaches.

### MGSE parameter selection and optimization

There are some MGSE parameter recommendations for optimal results. First, MGSE can accept different input types including both raw read files along with the reference assembly file, (sorted) BAM files, or coverage files. In many cases, BAM files are already available as a result of a typical genome assembly process which requires a read mapping to check the coverage of contigs as part of the quality control. Available BAM files avoid the computationally intensive step of read mapping as a part of MGSE’s execution, making the run more efficient.

Second, selection of the optimal reference regions for coverage calculation is crucial. Based on results obtained for *A. thaliana* and more complex genomes, it became evident that BUSCO genes work well as reference regions in a wide range of application scenarios. Hence, the preference of reference gene sets by the users can be BUSCO genes followed by all the genes. MGSE leverages established methods for coverage calculation by running bedtools thus no optimization is required for this process.

After choosing the regions for coverage calculation, it is necessary to filter out high coverage contigs that could distort the genome size estimations like contaminating sequences from microbes, mitochondrial sequences, and chloroplastic sequences. This blacklisting of high coverage contigs (>1.5x of average coverage) is activated by default, but can be deactivated by using the ‘--ignore’ flag. However, there are some cases in which it is not recommended to blacklist high coverage contigs, as shown for polyploids and organisms with a high degree of genomic repeats. Including the ‘--ignore’ flag can lead to better results in these cases. The user can run the busco2ploidy.py script to assess the apparent ploidy of the organism before running MGSE, in order to decide upon the optimal parameters. This script provides a frequency distribution plot of the BUSCO genes, and gives the user a ‘pseudo ploidy number’. Alternatively, the user can define the ‘--blacklist_factor’ to MGSE instead of using the ‘--ignore’ option. To use this method, the user can set the ‘--blacklist_factor’ to be ‘1.5 times the pseudo ploidy number’, which would heighten the cutoff for high coverage sequences, making it suitable for polyploid genomes and ensuring removal of just those sequences that exhibit unusually high coverage.

Further, coverage of the input datasets to MGSE significantly impacts the MGSE estimates. This is evident from the results of the coverage threshold analysis (AdditionalFile6). For *Arabidopsis*, the minimum number of bases (Mbp) required for optimal MGSE estimates from long reads was found to be 500 Mbp and the minimum number of bases required for optimal MGSE estimates from short reads was found to be 750 Mbp (Fig. 8a, 8b). While the minimum number of bases required for successful prediction by GenomeScope2 on short reads turned out to be 2500 Mbp (Fig. 8c). Given that the expected genome size of A. thaliana is ∼150 Mbp, the minimum coverage of datasets for a successful MGSE run based on long reads was found to be ∼3X, while that based on short reads was found to be ∼5X. These minimum coverage values were significantly lower than the minimum coverage of short read datasets needed for an optimal GenomeScope2 prediction, which was ∼17X. MGSE’s applicability to both long and short reads coupled with its ability to predict genome sizes based on datasets with low coverage values, makes it a valuable tool in addition to the existing kmer-based tools.

**Fig. 8:**
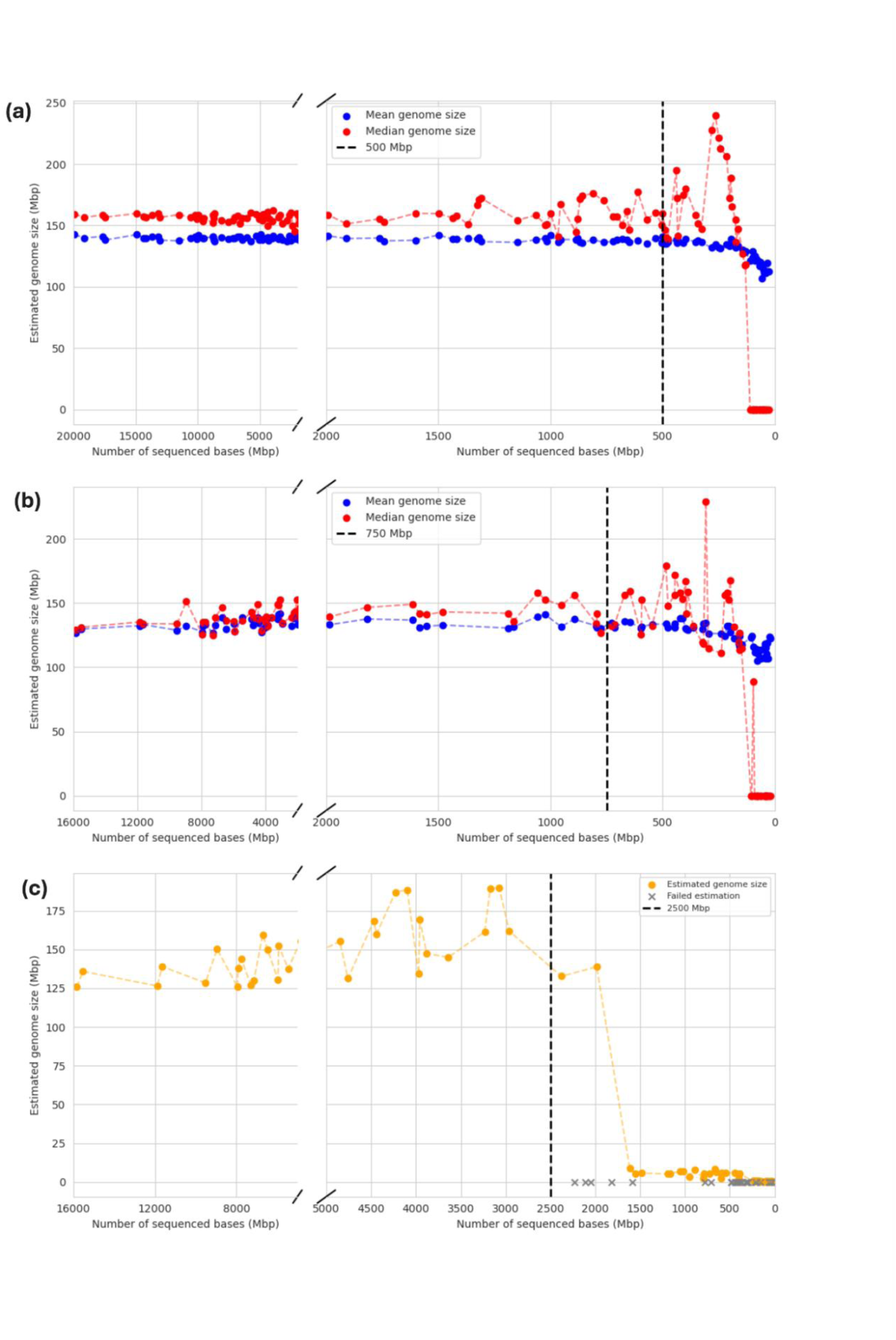
Coverage threshold analyses of (a) MGSE on long read datasets (b) MGSE on short read datasets (c) GenomeScope2 on short read datasets.

Finally, the runtime analyses of MGSE and GenomeScope2 revealed interesting patterns (AdditionalFile7). When comparing MGSE and GenomeScope2 in terms of the total runtime and the size of the input dataset, MGSE had higher runtimes than GenomeScope2 (Fig. 9). However, the computationally intensive step is the read mapping and MGSE could also be started with precomputed BAM files. When only considering the runtimes of coverage calculation and genome size prediction, the MGSE runtimes show a near plateau. For very large datasets, the MGSE and GenomeScope2 runtimes started to converge (Fig. 9). This indicates that MGSE can perform on par or even more efficiently than GenomeScope2 on very large datasets.

**Fig. 9:**
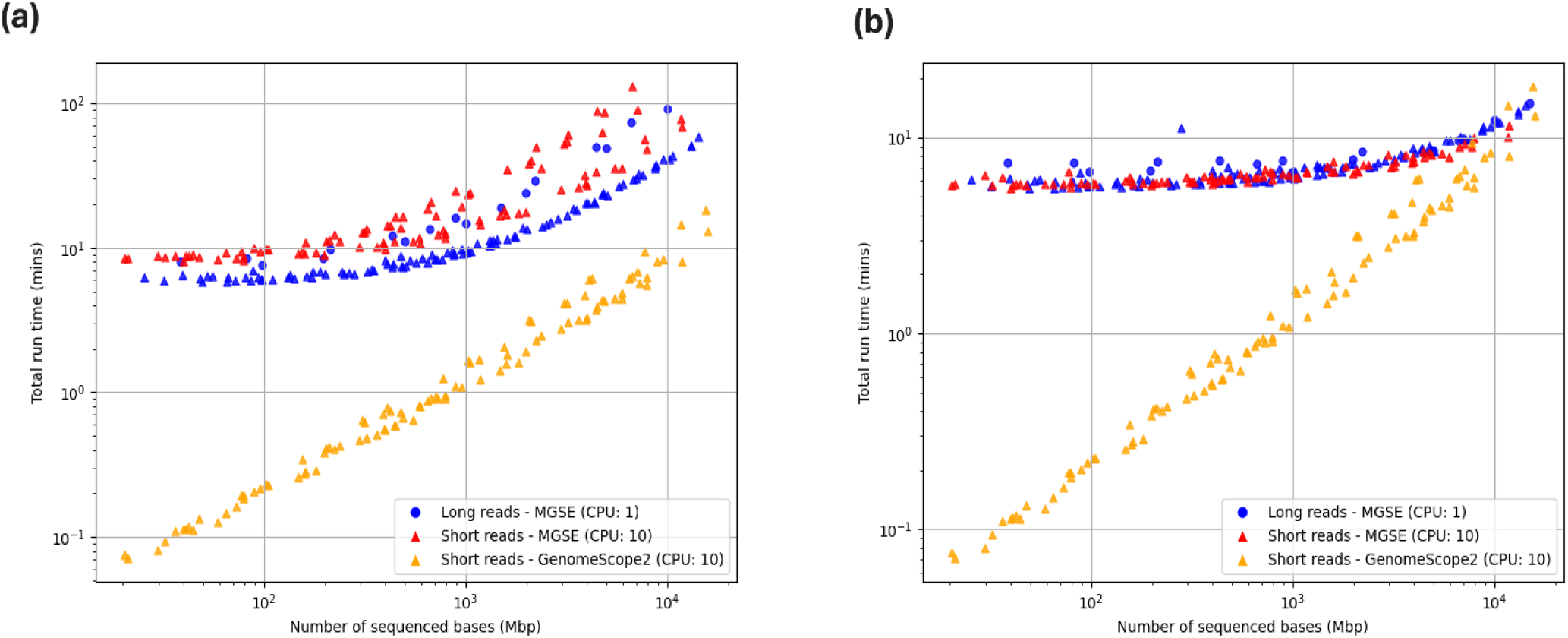
(a) Comparative analysis between total MGSE and GenomeScope2 runtimes and (b) Comparative analysis between coverage+MGSE execution times and GenomeScope2 total runtimes.

### Considerations about performance and outlook

MGSE performs best on a high contiguity assembly and requires a mapping of short or long reads to this assembly. Accurate coverage calculation for each position in the assembly is important and contigs display artificially low coverage values towards the ends. This is caused by a reduction in the number of possible ways in which reads can cover contig ends. The shorter a contig, the more the apparent coverage of this contig is reduced. Given that most assemblies are characterized by N50 values of >10Mbp, this effect can be neglected. Since a read mapping is required as input, MGSE might appear less convenient than classical k-mer-based approaches at first look. However, these input files are already available for many plant species, because such mappings are often part of the assembly (quality control) process [26, 27, 93, 94]. Future genome sequencing projects are likely to generate high continuity assemblies [25] and read mappings in the polishing process.

One advantage of MGSE is the possibility to exclude reads originating from contaminating DNA even if the proportion of such DNA is high. Unless reads from bacterial or fungal contaminations were assembled and included in the reference sequence, the approach can handle such reads without identifying them explicitly. This is achieved by discarding unmapped reads from the genome size estimation. MGSE works best with a high contiguity assembly and assumes all single copy regions of the genome are resolved and all repeats are represented by at least one copy. Although the amount of contamination of or in reads is usually small, such reads are frequently observed due to the high sensitivity of next generation sequencing methods [35, 95–97].

Reads originating from PCR duplicates could impact k-mer profiles and also predictions based on these profiles if not filtered out. After reads are mapped to a reference sequence, read pairs originating from PCR duplicates can be identified and removed based on identical start and end positions as well as identical sequences. This results in the genome size prediction by MGSE being independent of the library diversity. If the coverage is close to the read length or the length of sequenced fragments, reads originating from PCR duplicates cannot be distinguished from *bona fide* identical DNA fragments. Although MGSE results get more accurate with higher coverage, after exceeding an optimal coverage the removal of apparent PCR duplicates could become an issue. Thus, a substantially higher number of reads originating from PCR-free libraries could be used if duplicate removal is omitted. Depending on the sequencing library diversity, completely skipping the PCR duplicate removal step might be an option for further improvement. As long as these PCR duplicates are mapped equally across the genome, MGSE can tolerate these artifacts. This is in fact shown by the MGSE estimates of *E. coli*, *S. cerevisiae*, and *C. elegans* based on short read datasets. Removal of PCR duplicates from these datasets does not impact the MGSE size estimates (Fig. 7).

All methods are affected by DNA of the plastome and chondrome integrated into the nuclear chromosomes [98, 99]. K-mers originating from these sequences are probably ignored in many k-mer-based approaches, because they appear to originate from the chondrome or plastome, i.e., k-mers occur with very high frequencies. The apparent coverage in the mapping-based calculation is biased due to high numbers of reads which are erroneously mapped to these sequences instead of the plastome or chondrome sequence.

Differences in the GC content of genomic regions were previously reported to have an impact on the sequencing coverage [100, 101]. Both, extremely GC-rich and AT-rich fragments, respectively, are underrepresented in the sequencing output mainly due to biases introduced by PCR [102, 103]. Sophisticated methods were developed to correct coverage values based on the GC content of the underlying sequence [103–105]. The GC content of genes selected as reference regions for the coverage estimation is likely to be above the 36.3% average GC content of plants [77]. This becomes worse when only exons are selected due to the even higher proportion of coding sequence. Although a species-specific codon usage can lead to some variation, constraints of the genetic code determine a GC content of approximately 50% in coding regions. The selection of a large set of reference regions with a GC content close to the expected overall GC content of a genome would be ideal. However, the overall GC content is unknown as the GC content of regions not represented in the reference sequence is not known and cannot be inferred from the reads. As a result, the average sequencing coverage could be overestimated leading to an underestimation of the genome size. Future investigations would be necessary to develop a correction factor for this GC bias of short reads to further optimize the genome size prediction.

Many plant genomes pose an additional challenge due to recent polyploidy or high heterozygosity. Once high contiguity long read assemblies become available for these complex genomes, a mapping-based approach is feasible. As long as the different haplophases are properly resolved, the assessment of coverage values should reveal a good estimation of the genome size. Even the genomes of species which have recently undergone polyploidization could be investigated with moderate adjustments to the workflow. Reference regions need to be selected to reflect the degree of ploidy in their copy number.

With the widespread adoption of long read sequencing technologies in plant genomics, MGSE can turn out to be an important tool for genome size estimation. The major issue when developing tools for genome size prediction is the absence of a gold standard. In this study, predictions were compared against the best available genome sequences for the respective species. Several of these genome sequences should be very close to a perfect representation of the genome and thus have the potential to reveal the true genome size. Predictions generated by MGSE were constantly close to the sizes of these almost complete genome sequences. Moreover MGSE is able to handle low coverage datasets of both short and long reads, and still return reasonable genome size estimations. Finally, MGSE is universally applicable to all species and is not restricted to plants.

## Supporting information

AdditionalFile 1

AdditionalFile 2

AdditionalFile3

AdditionalFile4

AdditionalFile5

AdditionalFile6

AdditionalFile7

AdditionalFile8

AdditionalFile9

AdditionalFile10

AdditionalFile11

AdditionalFile12

AdditionalFile13

AdditionalFile14

## Data availability

Scripts developed as part of this work and a test data set are freely available on GitHub: https://github.com/bpucker/MGSE (https://doi.org/10.5281/zenodo.13984294). Underlying data sets are publicly available at the NCBI and SRA, respectively.

## Supplements

AdditionalFile1: Sequencing data set overview.

AdditionalFile2: *A. thaliana* genome size prediction values for all different approaches.

AdditionalFile3: *A. thaliana* genome size prediction of MGSE based on long reads

AdditionalFile4: *A. thaliana* genome size predictions by MGSE, findGSE, gce, and GenomeScope2.

AdditionalFile5: *B. vulgaris, Zea mays, Oryza sativa, Brachypodium distachyon, Solanum lycopersicum*, *Vitis vinifera, Fragaria x ananassa, Gossypium hirsutum, Eschericia coli, Saccharomyces cerevisiae,* and *Caenorhabditis elegans* genome size prediction values for all different approaches.

AdditionalFile6: Coverage threshold analyses on datasets for MGSE and GenomeScope2

AdditionalFile7: Runtime analyses of MGSE and GenomeScope2

AdditionalFile8: List of near complete/ complete and largest reported genome assembly sizes for the different organisms analyzed in this study

AdditionalFile9: Genome size estimation of *Brachypodium distachyon*.

AdditionalFile10: Genome size estimation of *Zea mays*.

AdditionalFile11: Genome size estimation of *Solanum lycopersicum*.

AdditionalFile12: Genome size estimation of *Vitis vinifera*.

AdditionalFile13: Genome size estimation of *Fragaria x ananassa*.

AdditionalFile14: Genome size estimation of *Gossypium hirsutum*.

## Author contributions

BP conceptualized the project, wrote the initial version of MGSE, performed several data analyses, generated figures, and wrote the initial manuscript version. SN updated the MGSE code, conducted additional data analyses, generated figures, and improved the manuscript. JG conducted additional data analyses and validated analyses. All authors read the final version of the manuscript and agreed to its submission.

## Conflict of interests

The authors declare that they have no conflict of interests.

## Acknowledgements

We are very grateful to the Bioinformatics Resource Facility support team of the CeBiTec for providing computing infrastructure and excellent technical support. Many thanks to members of the Genetics and Genomics of Plants (Bielefeld University) group who contributed to this work by discussion of preliminary results. Many thanks go to Hanna Schilbert, Nathanael Walker-Hale, and Iain Place for helpful comments on the manuscript. We also thank the Plant Biotechnology and Bioinformatics group (TU Braunschweig) for inspiring discussions. SN was supported by the doctoral scholarship of the German Academic Exchange Service (DAAD). This work was supported by the BMBF-funded de.NBI Cloud within the German Network for Bioinformatics Infrastructure (de.NBI) (031A532B, 031A533A, 031A533B, 031A534A, 031A535A, 031A537A, 031A537B, 031A537C, 031A537D, 031A538A).

## Notes

https://github.com/bpucker/MGSE

### Competing Interest Statement

The authors have declared no competing interest.

### Summary of Updates

- Improved the manuscript - Added coverage threshold and run time analyses - Added recommendations for optimal parameter choice - Demonstrated applicability beyond plants - Added script for ploidy inference

