## Supplementary figures and images for "Mapping-based Genome Size Estimation"

### AdditionalFile9

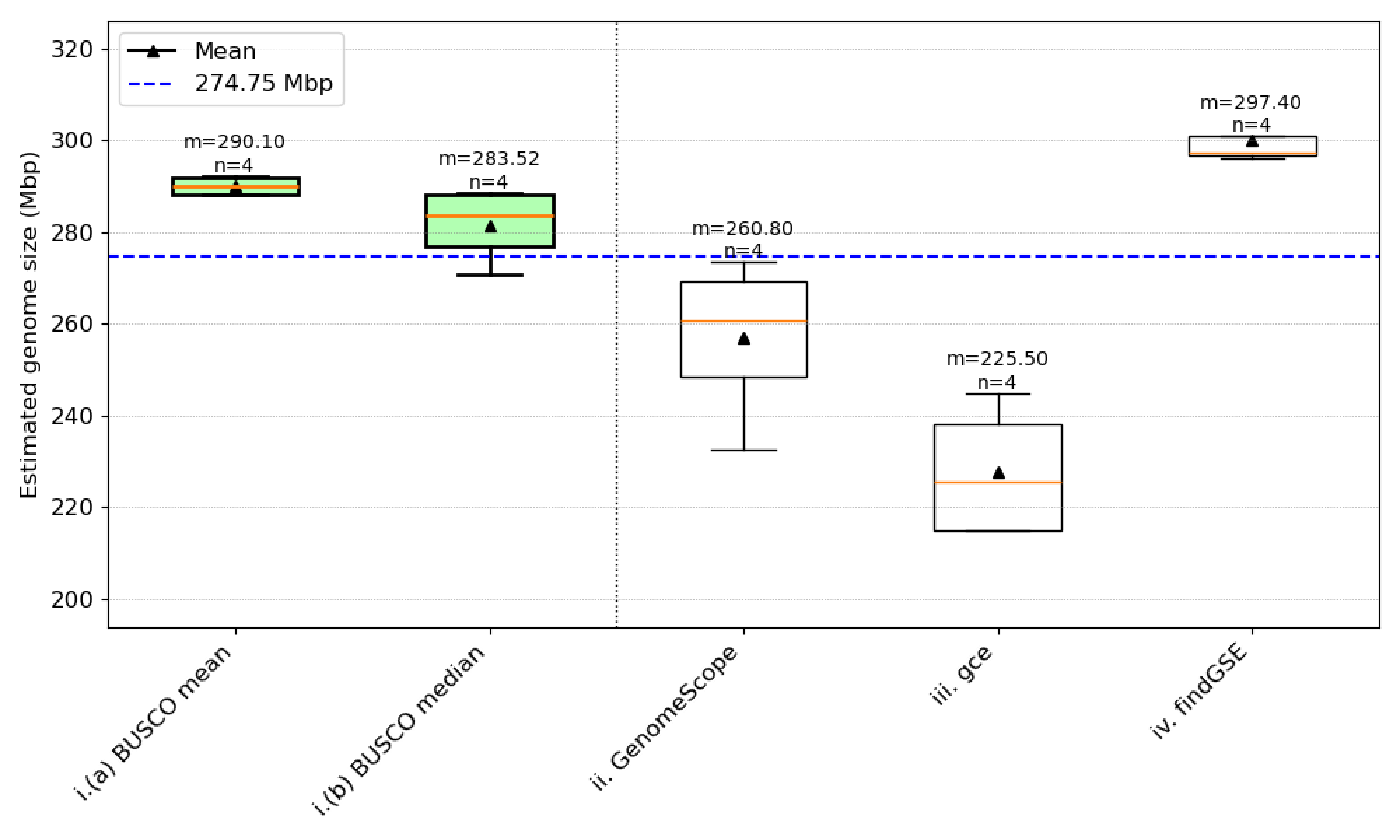

### AdditionalFile10

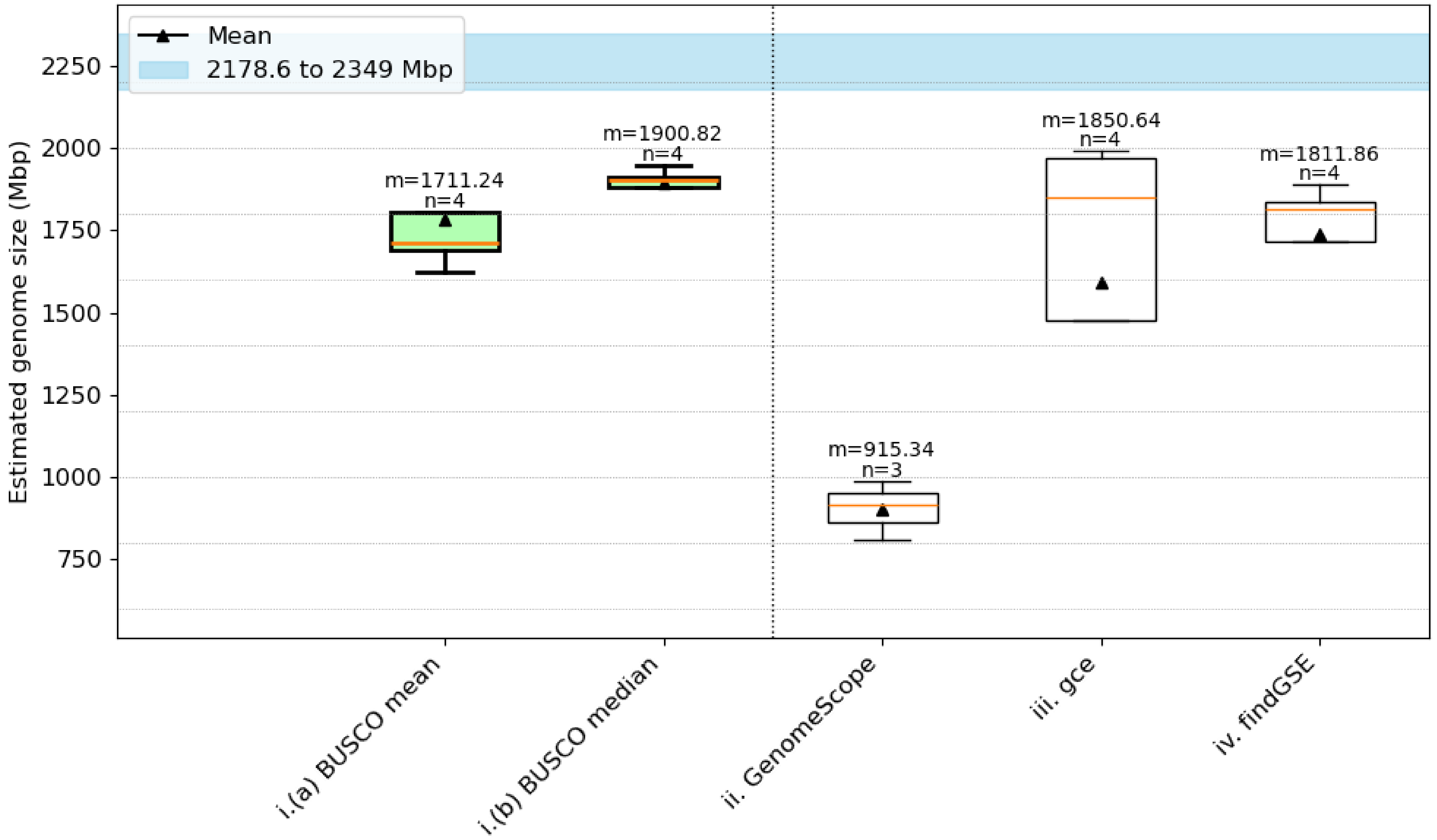

### AdditionalFile11

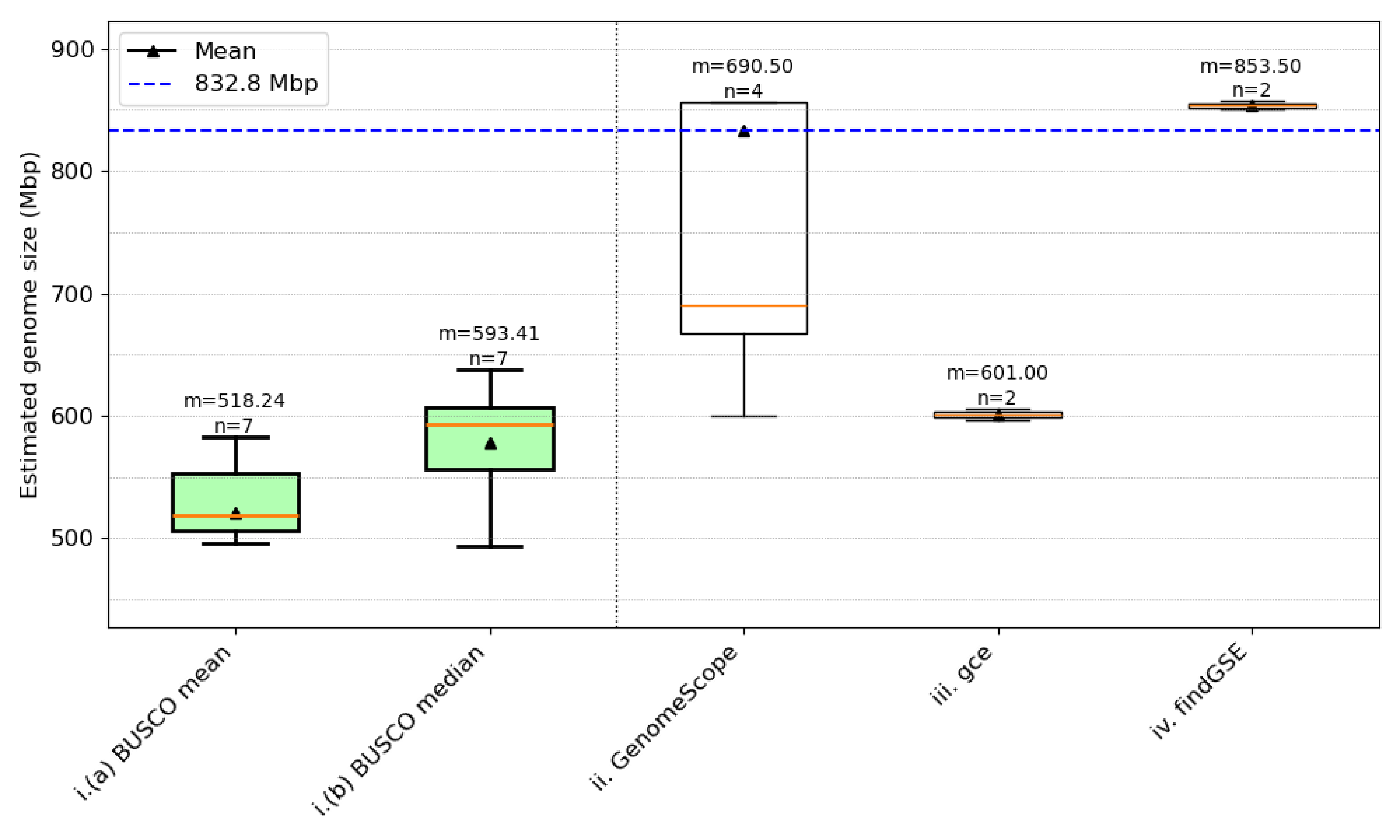

### AdditionalFile12

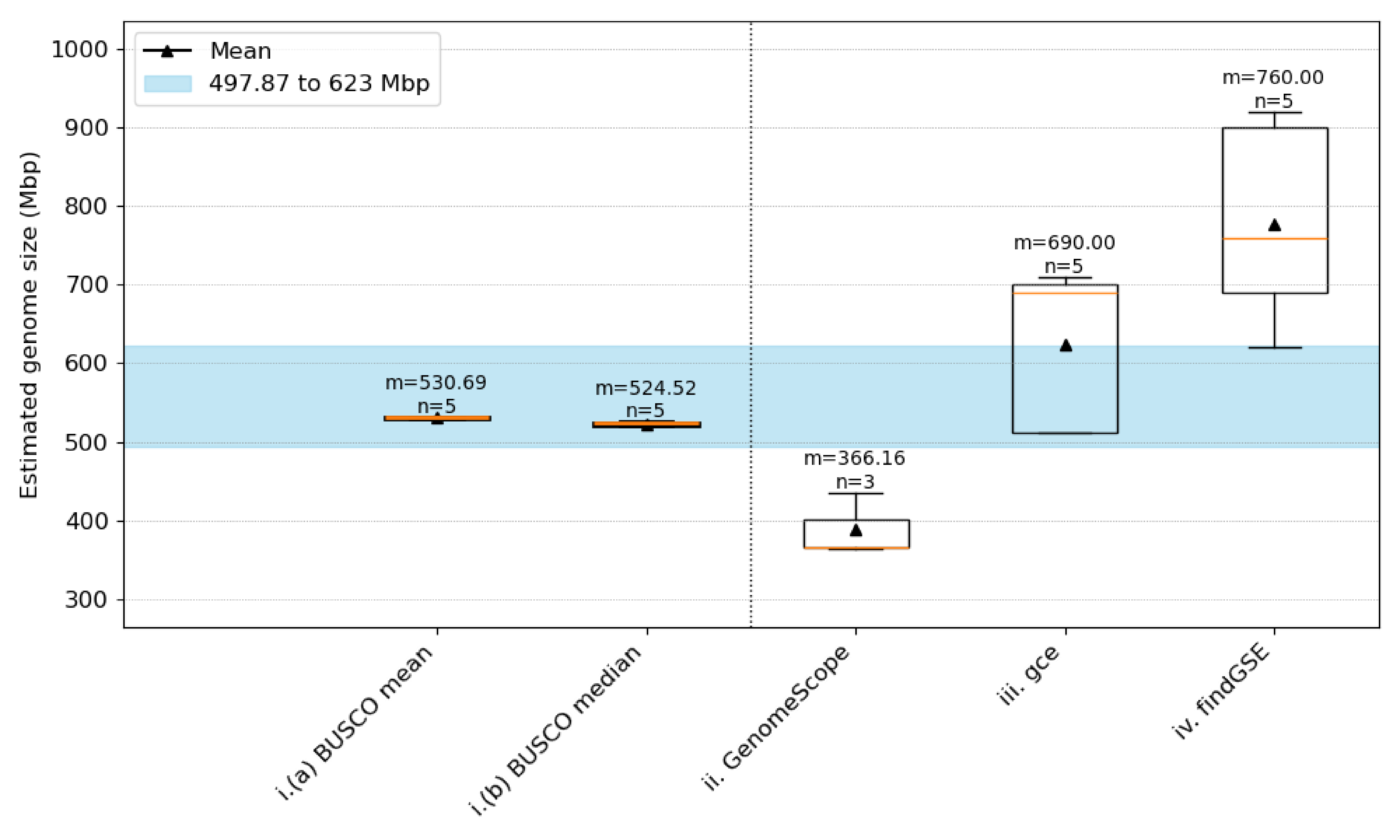

### AdditionalFile13

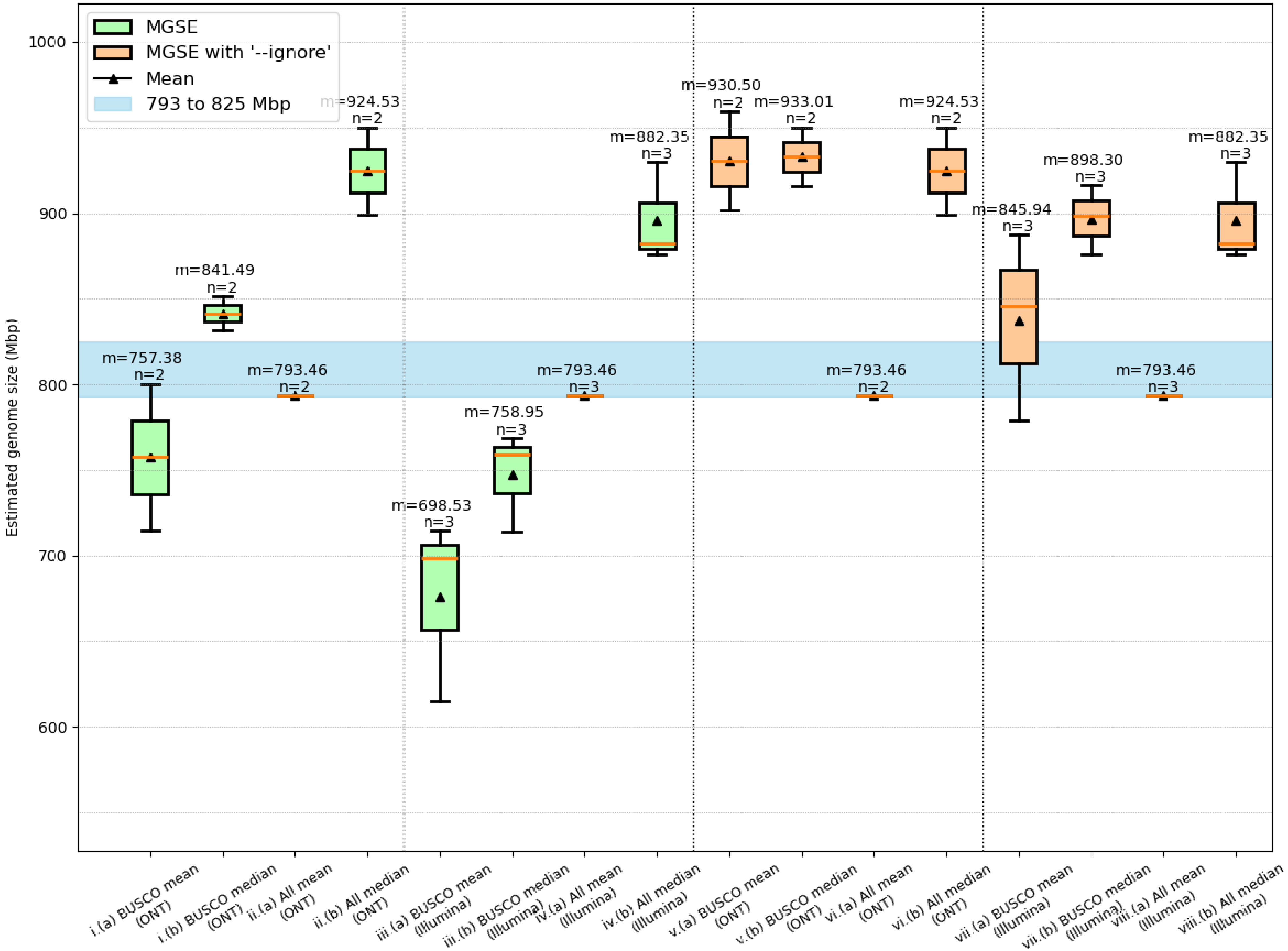

### AdditionalFile14

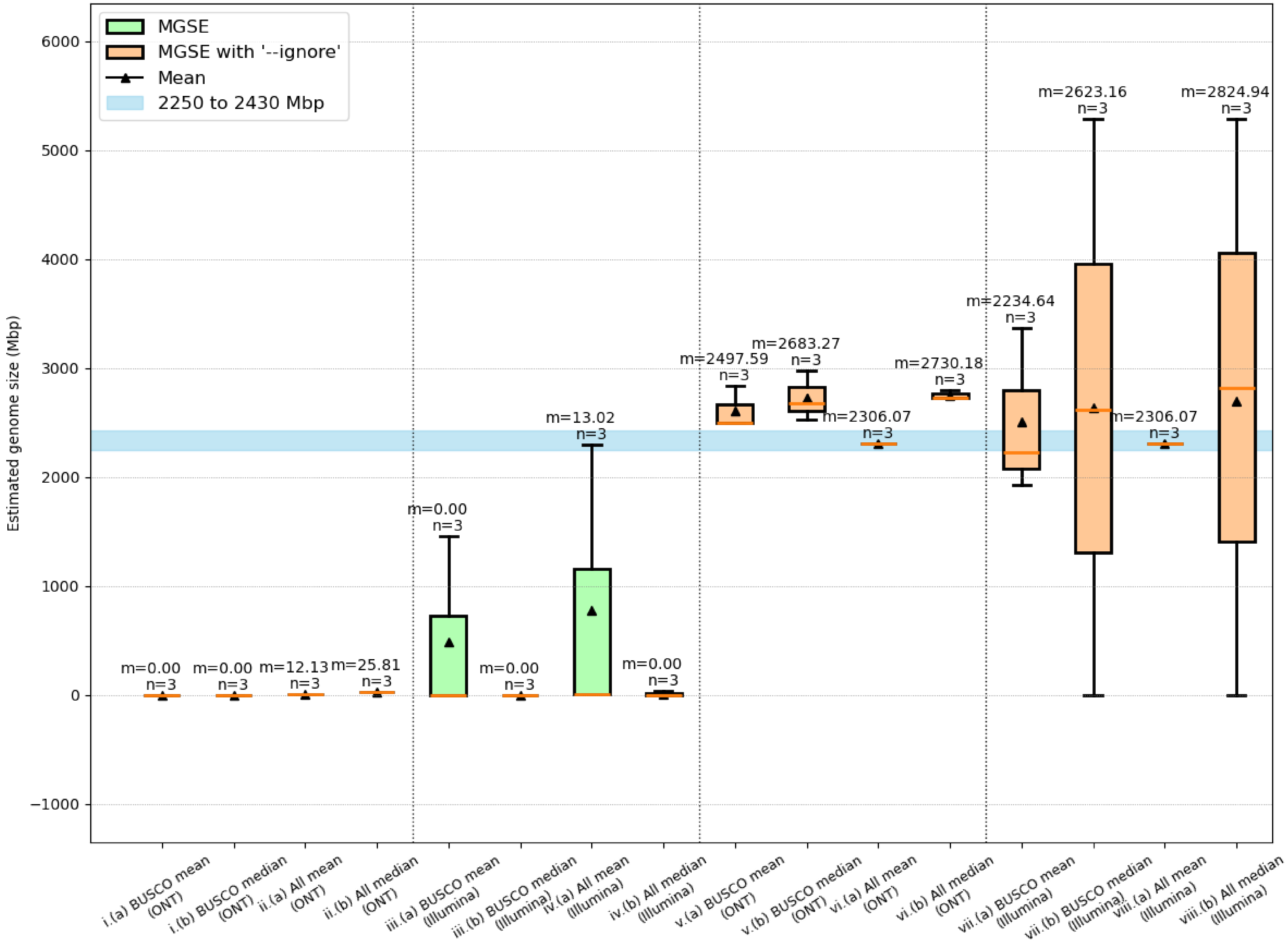
